# IFNγ induces epithelial reprogramming driving CXCL11-mediated T cell migration

**DOI:** 10.1101/2024.02.03.578580

**Authors:** Alessandro Cutilli, Suze A. Jansen, Francesca Paolucci, Michal Mokry, Enric Mocholi, Caroline A. Lindemans, Paul J. Coffer

**Affiliations:** Regenerative Medicine Center, University Medical Center Utrecht, Utrecht, The Netherlands; Center of Molecular Medicine, University Medical Center Utrecht, Utrecht, The Netherlands; Division of Pediatrics, University Medical Center Utrecht, Utrecht, The Netherlands; Princess Máxima Center for Pediatric Oncology, Utrecht, The Netherlands; Department of Cardiology, University Medical Center Utrecht, Utrecht, The Netherlands

## Abstract

The cytokine interferon-gamma (IFNγ) plays a multifaceted role in intestinal immune responses ranging from anti-to pro-inflammatory depending on the setting. Here, using a 3D co-culture system based on human intestinal epithelial organoids, we explore the capacity of IFNγ-exposure to reprogram intestinal epithelia and thereby directly modulate lymphocyte responses. IFNγ treatment of organoids led to transcriptional reprogramming, marked by a switch to a pro-inflammatory gene expression profile, including transcriptional upregulation of the chemokines CXCL9, CXCL10, and CXCL11. Proteomic analysis of organoid-conditioned medium post-treatment confirmed chemokine secretion. Furthermore, IFNγ-treatment of organoids led to enhanced T cell migration in a CXCL11-dependent manner without affecting T cell activation status. Taken together, our results suggest a specific role for CXCL11 in T cell recruitment that can be targeted to prevent T cell trafficking to the inflamed intestine.

## Introduction

The intestinal epithelium defines a physical barrier separating the intestinal lumen, where commensal and pathogenic microorganisms are located, and the internal tissues underneath. It is a monolayer tissue consisting of different secretory and absorptive intestinal epithelial cells (IECs) differentiating from intestinal stem cells (ISCs). The epithelium is organized in a villi-crypts architecture, with ISCs located at the bottom of the crypts^1^. The immune system of the gut is remarkably complex and consists of a variety of innate and adaptive immune cells that act to oppose a variety of possible threats^2^. Among other immune cells, T cells play a major role in several pathologies affecting the gastrointestinal tract, for instance, inflammatory bowel disease (IBD), coeliac disease, and acute intestinal graft-versus-host disease (GVHD)^2–9^.

Lymphocyte-mediated immune responses are elicited through trafficking of T cells to target tissues and establishment of local reactions. Indeed, these are features of all immune-related pathologies of the intestinal tract. IBD is an umbrella term that covers different pathologies including ulcerative colitis and Crohn’s disease. There is a distinct contribution of different T cell subsets for each disease, and this has recently been reviewed^10,11^. T cells also localize near intestinal crypts in early events leading to GVHD^12,13^, where they can release interferon-gamma (IFNγ), a cytokine at least in part responsible for the observed intestinal tissue damage resulting in enteric inflammation^14^. We have previously shown that IFNγ induces ISC death both *ex-vivo* and in murine models, compromising the regenerative potential of the intestinal epithelium^13^. Specifically, the stem cell compartment of the epithelium was protected in mice deficient for the IFNγ receptor (IFNγR)^13^. In coeliac disease patients, both CD4^+^ and CD8^+^ T cells are activated rapidly in response to gluten challenge^15^. They subsequently mediate an immune response characterized by IFNγ expression that correlates with intestinal tissue damage^16,17^. Similarly to GVHD, IFNγ-secreting T cells localize to the intestinal epithelium in coeliac disease^18^.

The role of IFNγ in pathogenic settings characterized by inflammation is complex due to its pleiotropic effects on different immune and non-immune cell types^19^. In IBD, IFNγ-expressing CD4^+^ T cells are positively associated with Crohn’s disease but not with ulcerative colitis, for which a negative association was found^20^. However, IFNγ disrupts the vascular barrier and has been showed to be inducing colitis in a murine experimental model^21^. In GVHD, donor derived IFNγ is protective at least in specific phases of disease pathogenesis. Depletion of donor CD4^+^ T cells shortly after allogeneic hematopoietic stem cell transplantation (allo-HSCT) results in higher serum levels of IFNγ, upregulation of PD-L1 in target tissues, and reduction of intestinal GVHD pathology^22^. Other studies have shown that lack of IFNγ induces lethal CD8^+^ T cell mediated-GVHD^23,24^, suggesting a potential protective effect. However, we have previously shown that allo-stimulated murine and activated human T cells reduce intestinal epithelial organoid viability ex-*vivo* in an IFNγ-dependent manner^13^. This effect was mediated through Janus kinase 1/2 (JAK1/2) signaling downstream of IFNγR^13^, and anti-IFNγ treated mice or mice treated with JAK1 inhibitors were protected from intestinal stem cell death.

Intestinal epithelial inflammation is the result of a systemic cascade, encompassing cellular activation, migration, and effector functionality. Given the complexity and the participation of diverse cell types across multiple tissues at distinct temporal stages, a comprehensive and precise delineation of interactions between T cells and IECs poses a challenge. A better understanding is required to expand the scope of potential therapeutic interventions in T cell-mediated intestinal disorders. The precise mechanism by which IFNγ shapes the inflammatory microenvironment in the gut and how it affects the crosstalk between ISCs, and T cells are still not completely clear. To address this, we have utilized an ex-*vivo* co-culture platform integrating human small intestinal organoids with peripheral blood-isolated T cells. Intestinal organoids are 3D epithelial cultures that originate from intestinal crypts when provided with appropriate growth factors and can differentiate into various types of IECs^25,26^. We and others have shown that organoids can serve as a pertinent ex-*vivo* model to examine interactions between intestinal epithelial cells and immune cells occurring *in vivo*^13,14,27–30^. Organoids treated with IFNγ show increased expression of inflammatory chemokines CXC motif chemokine ligand 9 (CXCL9), CXCL10 and CXCL11. Proteomic analysis confirmed the presence of those chemokines in the medium of IFNγ-exposed organoids. Using transwell migration assays and blocking antibodies we identified CXCL11 as the primary organoid-derived chemokine regulating T cell migration. Taken together, our results suggest a specific role for CXCL11 in T cell recruitment that can be explored for its therapeutic potential in preventing T cell trafficking to the inflamed intestine.

## Methods

### T cell isolation and activation

T cells were obtained from peripheral blood of healthy donors at UMC Utrecht, according to the METC approved protocol 07/125 or alternatively from buffy coats (Sanquin, NL). Following Ficoll-Paque (GE Healthcare) gradient separation, CD8^+^ T cells were isolated from the peripheral blood mononuclear cells (PBMCs) fraction in MACS buffer (2% heat-inactivated FBS, 2% 0,1M EDTA in PBSO) using the CD8^+^ Dynabead isolation kit (Thermo Fisher) and BD IMag Cell Separation Magnet (BD Biosciences). CD4^+^ T cells were isolated from the CD8-depleted PBMC fraction using the MagniSort human CD4^+^ T cell enrichment kit (Thermo Fisher). T cell purity was routinely checked by flow cytometry (>80%). T cells were activated using plate-bound functional grade anti-human CD3 (1.6 mg/ml in PBSO overnight at 4°C or 2h at 37°C, eBioscience) and soluble functional grade anti-human CD28 (1 mg/ml, eBioscience) for 3 or 4 days as indicated at a concentration of 1×10^6^ cells/ml in T cell medium (TCM) (RPMI Medium 1640+GlutaMAX-I, Gibco, with 100U/ml pen-strep and 10% heat-inactivated FBS).

### Intestinal organoid cultures

Healthy human small intestinal epithelial organoids were established and cultured as previously described^26^. In short, biobanked organoids were generated from biopsies of healthy individuals initially suspected of coeliac disease. All participants provided written informed consent as approved by the medical ethical review board of the UMC Utrecht (METC) (protocol METC 10-402/K; TCBio 19-489). Organoids (>passage 7) were passaged via single cell dissociation using 1x TrypLE Express (Gibco) or alternatively mechanically disrupted where indicated and resuspended in Advanced DMEM/F12 (Gibco), 100U/ml penicillin-streptomycin (Gibco), 10mM HEPES (Gibco) and Glutamax (Gibco) (GF-medium), and 50-66% Matrigel (Corning). After plating and Matrigel polymerization, human small intestinal organoid expansion medium (hSI-EM) was added consisting of GF-, Wnt-3a conditioned-medium (CM) (50%), R-spondin-1 CM (20%), Noggin CM (10%), murine EGF (50 ng/ml, Peprotech), nicotinamide (10 mM, Sigma), N-acetyl cysteine (1.25 mM, Sigma), B27 (Gibco), TGF-β inhibitor A83-01 (500 nM, Tocris), p38 inhibitor SB202190 (10 µM, Sigma), Rho-kinase/ROCK inhibitor Y-27632 (10 µM, Abcam, for the first 2-3 days of culture), and Primocin (optional) (100 µg/ml, Invitrogen). Medium was changed every 2-3 days. For indicated timepoints, treatment wells received 10 ng/ml (for migration assays, co-culture assays proteomic screening) and or 1 ng/ml (for RNA-seq) of rhIFNγ (R&D systems).

### Migration transwell assay

Organoids were cultured in 24-well plates and treated for 48h with indicated conditions. After IFNγ treatment (10 ng/ml), the medium was changed with fresh hSI-EM without p38 inhibitor (no SB) for 24h. In the meantime, T cells were activated for 3 days or left resting in TCM in preparation of the assay. At the start of the assay, the T cells were stained with CellTrace Violet (CTV) and added to 3 µm-pored transwell inserts (Greiner Bio-One) (4×10^5^ T cells in 200 µl) that were inserted in the wells containing organoids. After overnight incubation, inserts were removed, and the contents of each well was dissociated with TrypLE and reconstituted in 300 µl. The number of CTV^+^ events per 150 µl sample was counted using flow cytometry. For the chemokine blocking assays, 1 µg/ml anti-CXCL9, anti-CXCL10 or anti-CXCL11 mAb (R&D systems) were added to the lower compartment prior to start of the assay.

### T cell activation co-culture assay

For the measurement of T cell activation and proliferation in the presence of IFNγ-treated organoids, organoids were cultured and treated with IFNγ (10 ng/ml) for 24h. At the end of the incubation time, organoids were mechanically disrupted. A portion of each condition was used to dissociate into single cells to infer the cell number and normalize the split ratios between treated condition and untreated controls. with cells were stained with CTV prior of co-culture. Co-cultures were plated in a 96-well plate, in a final volume of 200 µl containing 2×10^5^ T cells and the equivalent of 1/6^th^ of untreated organoids per well, in no SB medium with 10% BME. To activate T cells, wells were previously coated with anti-CD3 and the culture medium was supplemented with anti-CD28. After 4 days of co-culture, plates were cooled at 4°C for 30 min to dissolve BME and centrifuged at 500g 5 min at 4°C. For intracellular staining, GolgiStop (containing Monensin, BD Biosciences) was added to the medium 4h before collecting cells for the staining. The content of each well was dissociated to single cells with TrypLE, washed with PBSO and consequently stained as described in the Flow Cytometry section.

### Imaging of organoids

Bright field (culture) images were acquired using an EVOS FL Cell Imaging System (Thermo Fisher Scientific).

### Flow Cytometry

T cells were stained with live/dead marker Zombie NIR (BioLegend) and fluorochrome-labeled antibodies anti-CD3-PE and anti-CD4- or anti-CD8-FITC (BioLegend) either in FACS buffer (PBSO, 2mM EDTA, 0.5% BSA, Sigma) or MACS buffer (PBS0, 2mM EDTA, 2% FBS). For measuring T cell activation, anti-CD25-APC (BioLegend), anti-CD69-BV605 (BioLegend) were added to the staining. Intracellular IFNγ-staining was performed using the Intracellular Fixation & Permeabilization Buffer Set (eBioscience Thermofisher) with anti-IFNg-PECy7 (BD Biosciences). For measuring proliferation, T cells were stained before the start of the assay with CellTraceViolet (CTV) (Invitrogen, 5 µM in PBS0) according to manufacturer’s protocol. For assessing CXCR3 and CXCR7 expression, anti-CXCR3-PE and anti-CXCR7-APC (BioLegend). FC data were acquired with a BD LSRFortessa Cell Analyzer (BD Biosciences) using FACSDiva (BD Biosciences) software and a CytoFLEX Flow Cytometer (Beckman Coulter) with CytExpert software. The data were analyzed with FlowJo (Treestar, 10.6.2) or CytExpert software (2.4).

### RNA sequencing

For RNA sequencing, mRNA from organoids treated with IFNγ (1 ng/ml) was isolated using Poly(A) Beads (NEXTflex). Sequencing libraries were prepared using the Rapid Directional RNA-Seq Kit (NEXTflex) and sequenced on a NextSeq500 (Illumina) generating 75 base long reads (Utrecht DNA Sequencing Facility). The obtained reads were mapped against the reference genome (hg19 assembly, NCBI37) using BWA41 package (mem –t 7 –c 100 –M –R)^31^. Analysis was performed using DESeq2^32^ in R2: Genomics Analysis and Visualization Platform (http://r2.amc.nl). Analysis consisted of principal component analysis (PCA), and generation of the list of differentially expressed genes (DEGs) (padj<0.05). Gene Ontology (GO) term analysis was performed using either upregulated (log2FoldChange>0) or downregulated (log2FoldChange<0) DEGs as compared to control per condition using 2X2 contingency table analysis chi-square with continuity correction (padj<0.05). REVIGO software was used to visualize GO term clusters (http://revigo.irb.hr/). Gene Set Enrichment Analysis Pre-ranked analysis was performed with the GSEA software checking for enrichment of genes part of Hallmark datasets in the GSEA software^33^.

The dataset analyzed in this manuscript is uploaded on EGA (EGA ID: EGAS50000000083).

### Olink proximity extension proteomic analyses

Organoids were cultured in 24-well plates and treated for 48h with IFNγ 10 ng/ml. After treatment, the medium was changed with fresh hSI-EM without p38 inhibitor (no SB) for 24h. At the end of the incubation, the CM was collected and centrifuged 1000g for 5 min. The supernatant was transferred to new Eppendorf tubes and stored at -80 °C until further processing. The Olink Target 96 Immuno-Oncology panel (v.3112) from Olink (Uppsala, Sweden) was used to quantify 92 immuno-oncology related proteins in each sample (IL-8, TNFRSF9, TIE2, MCP-3, CD40-L, IL-1 alpha, CD244, EGF, ANGPT1, IL-7, PGF, IL-6, ADGRG1, MCP-1, CRTAM, CXCL11, MCP-4, TRAIL, FGF2, CXCL9, CD8A, CAIX, MUC-16, ADA, CD4, NOS3, IL-2, Gal-9, VEGFR-2, CD40, IL-18, GZMH, KIR3DL1, LAP TGF-beta-1, CXCL1, TNFSF14, IL-33, TWEAK, PDGF subunit B, PDCD1, FASLG, CD28, CCL19, MCP-2, CCL4, IL-15, Gal-1, PD-L1, CD27, CXCL5, IL-5, HGF, GZMA, HO-1, CXCL1, CXCL10, CD70, IL-10, TNFRSF12A, CCL23, CD5, CCL3, MMP7, ARG1, NCR1, DCN, TNFRSF21, TNFRSF4, MIC-A/B, CCL17, ANGPT2, PTN, CXCL12, IFN-gamma, LAMP3, CASP-8, ICOSLG, MMP12, CXCL13, PD-L2, VEGFA, IL-4, LAG3, IL12RB1, IL-13, CCL20, TNF, KLRD1, GZMB, CD83, IL-12, CSF-1). Multiplex proximity extension assay panels were used to quantify each protein, as previously described^34^. The raw quantification cycle values were normalized and converted into normalized protein expression (NPX) units. The NPX values were presented on a log2 scale in which one unit higher in NPX values represents a doubling of the measured concentration. The quality control of the measured samples was conducted following Olink’s standard quality control protocol.

### Statistical analysis

Data are presented as mean ± standard error of the mean (SEM). To take into account intra-individual and intra-experimental variation experiments were performed at least twice, and sample material coming from at least two different human donors. Statistical significance was determined at *P ≤* 0.05 using one-way analysis of row-matched (RM) variance (ANOVA) with Dunnett’s multiple comparison test or a Student *t* test where appropriate. Significance is indicated as *P ≤* 0.05 (*), *P ≤* 0.01 (**), or *P ≤* 0.001 (***) or *P* < 0.0001 (****).

### Data sharing statement

For original data please contact

## Results

### IFNγ induces intestinal epithelial transcriptional reprogramming leading to CXCL9, CXCL10 and CXCL11 release

To study the effect of IFNγ-exposure on IECs, we first treated human small intestinal organoids with IFNγ for 48 hours. IFNγ-exposure resulted in visible morphological changes, smaller organoid size with thickened membrane, a condensed phenotype with shedding of dead cells, and cell debris inward into the organoid lumen (**Suppl. Figure 1a**). To evaluate transcriptional responses to IFNγ-treatment, bulk RNA sequencing (RNA-seq) of three independent organoid lines was performed. Exposure to IFNγ resulted in gene expression changes 6h and 18h after treatment (**Figure 1a**). Notably, untreated control organoids manifested a more substantial separation along PC1 between different organoid lines compared to the IFNγ-treated organoids. This suggests that IFNγ-induced reprogramming results in robust responses across different genetic and transcriptional baselines of three donors. IFNγ treatment resulted in 448 DEGs after 6h treatment and in 754 DEGs after 18h treatment (**Figure 1b, Suppl. Figure 1b**, **Supp. Table 1**). A list of the top 50 DEGs (25 most upregulated and 25 most downregulated genes) identified for each treatment duration is shown in **Figure 1c** and **d** with a visual representation of their log2 fold change (log2FC). In general, the 386 DEGs shared by the two treatment durations show a more pronounced expression change over time (**Figure 1e**). Chemokines CXCL9, CXCL10 and CXCL11 were rapidly and strongly upregulated: *CXCL9* 10.78 log2FC at 6h and 11.48 log2FC at 18h, *CXCL10* 11.52 log2FC at 6h and 11.42 log2FC at 18h, *CXCL11* 9.78 log2FC at 6h and 9.93 log2FC at 18h. GO term analysis of DEGs indicated an over-representation of genes belonging to gene sets including “immune response”, “response to interferon-gamma”, “cell-cell junction organization”, and “response to transforming growth factor beta” (**Figure 1f**; a visual representation of significant GO gene sets is provided in **Suppl. Figure 1c**; a complete list of GO gene sets and related DEGs is provided in **Suppl. Table 1** for 6h and 18h treatment). Furthermore, GSEA showed not only a positive correlation of IFNγ-regulated genes with those previously associated with IFNγ-responses and JAK-STAT signaling, but also with inflammatory responses and allograft rejection supporting the biological significance of our observations (**Figure 1g**).

**Figure 1.**
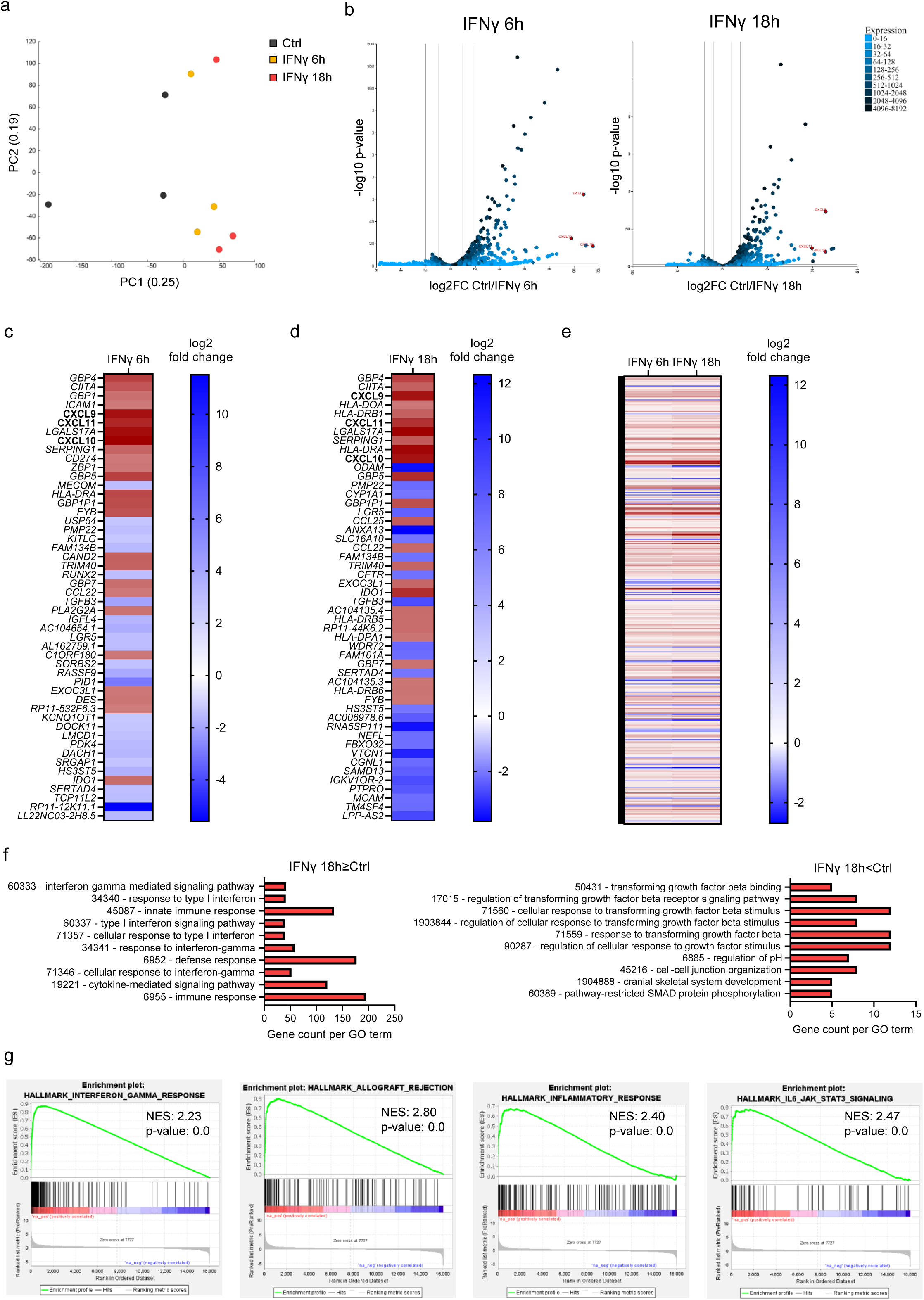
IFNγ exposure reprograms the intestinal epithelial transcriptome. Human small intestinal epithelial organoids (n=3 organoid lines) were treated with IFNγ (1 ng/ml) for 6h and 18h. RNA was isolated and subjected to bulk RNA-sequencing, after which bioinformatics analyses was performed. **(a)** PCA clustering of samples from three organoid lines used to perform RNA-seq after no treatment, IFNγ (1 ng/ml) for 6h and IFNγ (1 ng/ml) for 18h. Each point represents an individual sample from an organoid line, color-coded for the treatment and exposure time. **(b)** Volcano plots indicating differentially expressed (padj<0.05) genes of 6h (left) and 18h (right) IFNγ-treated organoids versus untreated control organoids. *CXCL9*, *CXCL10*, and *CXCL11* are annotated. (**c**) Heatmaps of top 50 most different differentially expressed genes (DEGs) (25 top upregulated, 25 top downregulated) after treatment with IFNγ for 6h compared to untreated controls (DESeq2, padj<0.05). DEGs are ordered by p-value, with the most significant genes in the upper part of the list. **(d)** Heatmaps of top 50 most DEGs (25 top upregulated, 25 top downregulated) after treatment with IFNγ for 18h compared to untreated controls (DESeq2, padj<0.05). DEGs are ordered by p-value, with the most significant genes in the upper part of the list. **(e)** Heatmaps of all significant DEGs shared by the conditions in which organoids have been treated with IFNγ for 6h and 18h compared to untreated controls (DESeq2, padj<0.05). DEGs are ordered alphabetically from A to Z. **(f)** Most significant Gene Ontology (GO) term analysis of Biological Processes in genes upregulated (LFC>0) or downregulated (LFC<0) by treatment with IFNγ for 18h compared to untreated controls (pdaj<0.05). **(e)** Gene Set Enrichment Analysis (GSEA) on all genes in organoids treated with IFNγ for 18h compared to untreated controls.

To determine whether the observed intestinal organoid transcriptional reprogramming also resulted in changes in protein secretion we employed Olink proteomics analysis. Organoids were treated for 48h with IFNγ followed by washing and an additional 24h of culture in IFNγ-free fresh medium. The conditioned media (CM) from untreated control organoids and IFNγ-treated organoids from three lines were evaluated by the Olink proteomics platform. 44 proteins were above the level of detection (LOD) (**Figure 2a**), with significant changes in the levels of CXCL9, CXCL10, and CXCL11 (**Figure 2b**), in line with the high epithelial expression of these genes.

**Figure 2.**
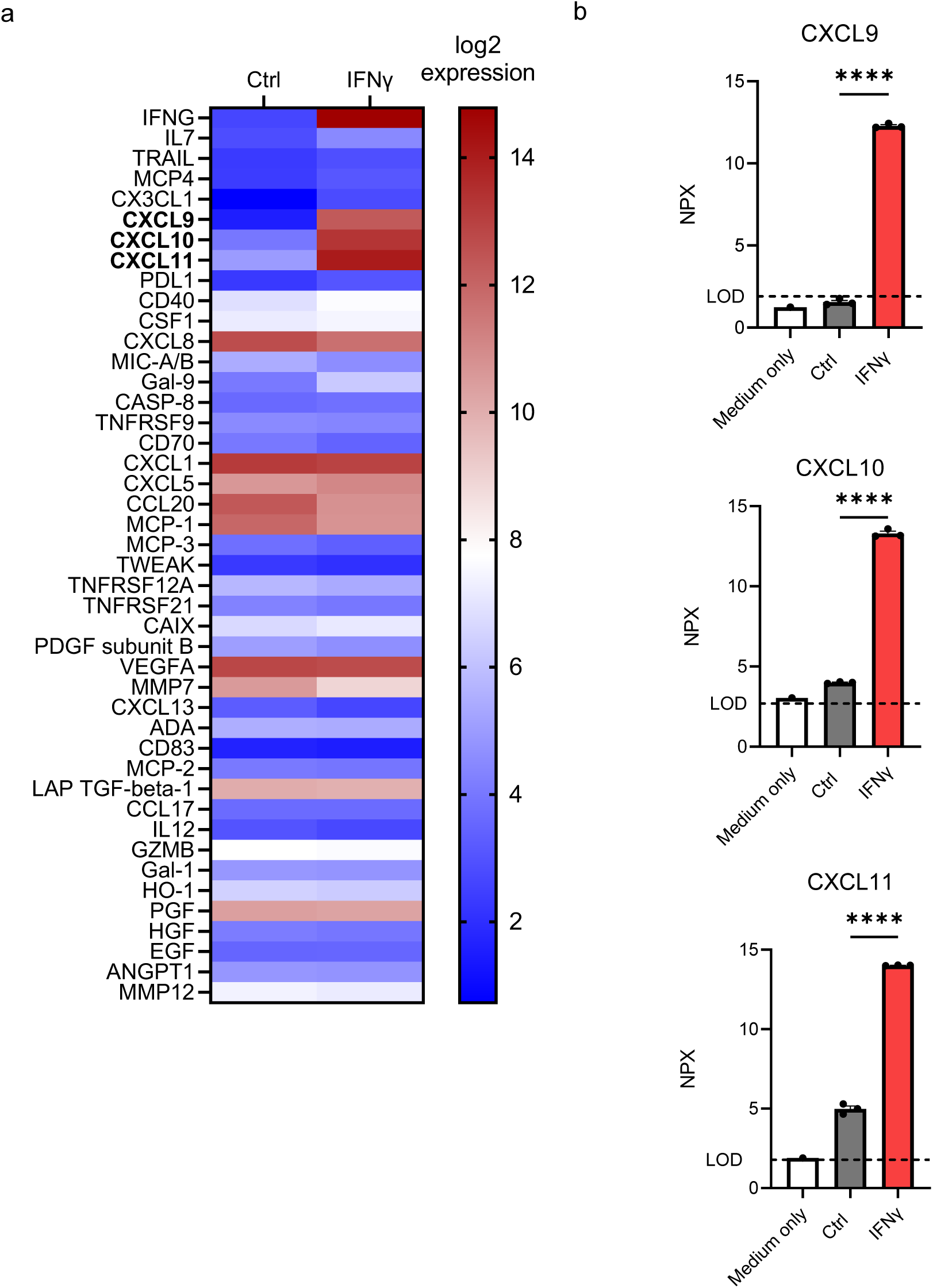
IFNγ-treatment increases expression and release of CXCL9, CXCL10, and CXCL11 from intestinal organoids. **(a)** Analysis of proteins present in conditioned media of organoids treated with IFNγ (10 ng/ml) for 48h (Olink Proteomics) (n=3 donors). **(b)** CXCL9, CXCL10, and CXCL11 levels detected by Olink proteomics. Each data point indicates the levels of the detected protein in the CM from each organoid condition (log2 scale), mean with SEM, ANOVA.

Taken together, these data demonstrate that human small intestinal organoids respond to IFNγ by acquiring a pro-inflammatory gene expression profile, characterized by expression and secretion of CXCL9, CXCL10, and CXCL11.

### IFNγ-induced intestinal epithelial damage enhances T cell migration

Since IFNγ induced chemokine expression in intestinal epithelia we aimed to evaluate whether IFNγ-treated organoids can directly influence T cell responses, specifically migration. We made use of a transwell system with 3 μm-pored inserts. Organoids were treated with IFNγ for 48 hours and subsequently cultured in IFNγ-free medium for an additional 24 hours. Either unstimulated or polyclonally pre-activated CTV-stained human peripheral blood CD4^+^ and CD8^+^ T cells were subsequently added to the transwell insert. After 24 hours, the number of T cells that had migrated to the lower (organoid) compartment was evaluated. Both unstimulated and pre-activated CD4^+^ and CD8^+^ T cells demonstrated significantly increased migration towards IFNγ-treated organoids (**Figure 3a-b**).

**Figure 3.**
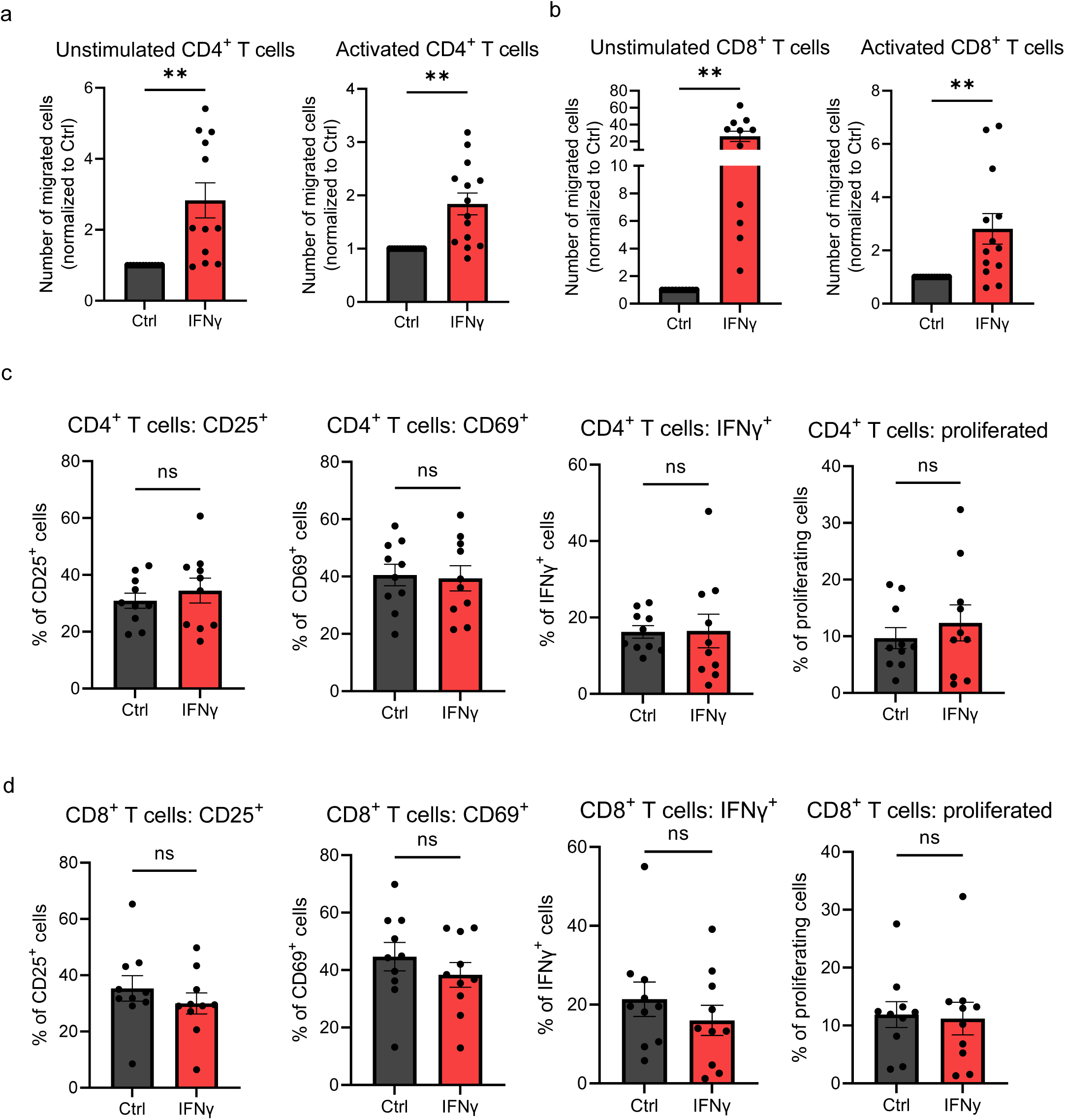
IFNγ-induced intestinal epithelial damage enhances T cell migration but does not alter T cell activation and proliferation. **(a-b)** Normalized number of unstimulated and pre-activated CD4^+^ and CD8^+^ T cells that have migrated overnight from a 3 μm-pore sized insert (upper compartment) to the lower compartment containing organoids that were treated with IFNγ (10 ng/ml) for 48h, and then refreshed for 24h prior to the beginning of the assay, counted by FACS. n≥11 T cell donors with 1 organoid line, each data point indicates a T cell donor, mean with SEM, Student *t* test. **(c-d)** (Membrane) activation markers CD25 and CD69 expression, IFNγ intracellular expression and proliferation of CD4^+^ and CD8^+^ T cells as measured by CTV-dilution after 4-day co-culture with organoids that were previously treated with IFNγ (10 ng/ml) for 24h before the start of the assay. Organoids were washed and disrupted mechanically before replating in co-culture with T cells. n≥10 T cell donors with 1 organoid line, each data point indicates a T cell donor, mean with SEM, Student *t* test.

Next, the effect of IFNγ-damaged epithelium on T cell activation was investigated. CD4^+^ and CD8^+^ T cells were isolated, stained with CTV, and activated by incubation with plate-bound anti-CD3 and soluble anti-CD28, in the presence of untreated or IFNγ-treated organoids. Proliferation and activation markers were assessed by flow cytometry after four days of co-culture. Membrane expression of activation markers CD25 and CD69, as well as IFNγ production and proliferation, were unaltered between T cells co-cultured with untreated control organoids or pre-treated organoids (**Figure 3c-d**).

In summary, IFNγ-treatment of intestinal organoids specifically increased T cell migration, without having an additional effect on T cell activation.

### CXCL11, but not CXCL9 and CXCL10, modulates T cell migration towards IFNγ-treated organoids

To determine the relative contribution of organoid-derived CXCL9, CXCL10 and CXCL11 on T cell migration, chemokine blocking assays were performed. Specific monoclonal antibodies (mAbs) with reported neutralizing activity were added to the lower compartment of the transwell system for the duration of the assay. In conditions where untreated control organoids were present, there was no significant effect on unstimulated CD4^+^ and CD8^+^ T cell migration when blocking antibodies were utilized, while we observed a significant reduction of pre-activated T cell migration towards organoids when anti-CXCL11 was used (**Figure 4a-b**). When organoids were previously treated with IFNγ, blocking CXCL11 resulted in decreased migration for both unstimulated and pre-activated CD4^+^ and CD8^+^ T cells (**Figure 4c-d**). In contrast to CXCL11, blocking CXCL9 and CXCL10 had no effect on T cell migration towards organoids.

**Figure 4.**
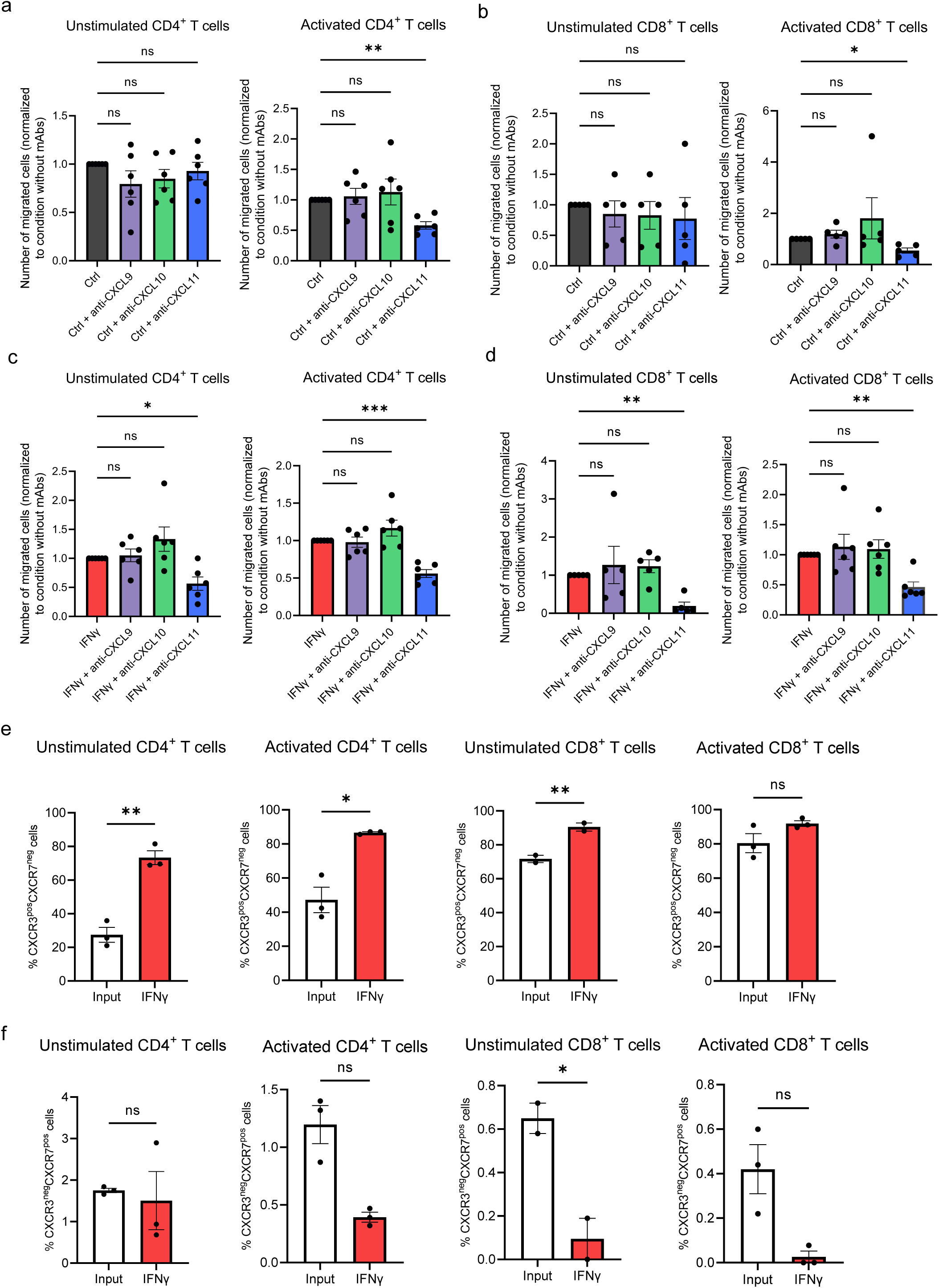
CXCL11, but not CXCL9 and CXCL10, modulates T cell migration towards IFNγ-treated organoids. **(a-b)** Normalized number of unstimulated and pre-activated CD4^+^ and CD8^+^ T cells that have migrated overnight from a 3 μm-pore sized insert (upper compartment) to the lower compartment containing untreated control organoids, counted by FACS. mAbs with neutralizing activity have been used to block CXCL9, CXCL10, or CXCL11 (1 µg/ml). n≥5 T cell donors with 1 organoid line, each data point indicates a T cell donor, mean with SEM, ANOVA. **(c-d)** Normalized number of unstimulated and pre-activated CD4^+^ and CD8^+^ T cells that have migrated overnight from a 3 μm-pore sized insert (upper compartment) to the lower compartment containing organoids that were treated with IFNγ (10 ng/ml) for 48h, and then refreshed for 24h, counted by FACS. mAbs with neutralizing activity have been used to block CXCL9, CXCL10, or CXCL11 (1 µg/ml). n≥5 T cell donors with 1 organoid line, each data point indicates a T cell donor, mean with SEM, ANOVA. **(e-f)** (Membrane) CXCR3 and CXCR7 expression of T cells that have migrated overnight from a 3 μm-pore sized insert (upper compartment) to the lower compartment containing organoids that were treated with IFNγ (10 ng/mL) for 48h, and then refreshed for 24h prior to the beginning of the assay. % of positive CXCR3 single positive cells **(e)** or CXCR7 single positive cells **(f)** are compared to the percentage in the input material, which consists of T cells cultured for the whole duration of the migration assay in a well containing empty Matrigel drops (no organoids). N≥2 T cell donors with 1 organoid line, each data point indicates a T cell donor, mean with SEM, Student *t* test.

We observed higher levels of IFNγ in the CM from IFNγ-treated organoids despite the medium change (**Figure 2a**, **Suppl. Figure 2a**). To evaluate whether residual IFNγ might be implicated in the observed effects on T cell migration, we performed migration assays with empty matrigel drops, subjected to the same processing as the other samples containing organoids. We observed no differences in the migration of T cells towards matrigel either cultured in IFNγ-containing medium or IFNγ-free medium, confirming that possible residual IFNγ is not influencing T cell migration (**Suppl. Figure 2b**). These results suggest that epithelial-organoid-expressed CXCL11 specifically recruits T cells to IFNγ-treated intestinal epithelium.

CXCL9, CXCL10 and CXCL11 share the same receptor, CXCR3^35^. In addition, CXCL11 has also been reported to bind CXCR7^36^. To determine which chemokine receptor was involved in T cell migration to IFNγ-treated organoids, T cells were collected from the lower compartment of the transwell system and CXCR3 and CXCR7 expression levels were analyzed with flow cytometry. T cells migrating towards IFNγ-treated organoids were enriched for CXCR3 expression compared to input, which consisted of the cells used in the migration assay kept at culture conditions with empty matrigel drops and processed alongside the other samples (**Figure 4e**). The only exception found was related to pre-activated CD8^+^ T cells, of which a high percentage of cells were already expressing CXCR3 in the input condition. The percentage of CXCR7-expressing T cells was negligible, considering either its expression alone (**Figure 4f**) or in combination with CXCR3 (**Suppl. Figure 2c**).

Taken together, our results demonstrate that CXCL11 released from IFNγ-exposed intestinal epithelium promotes both CXCR3^+^CD4^+^ and CXCR3^+^CD8^+^ T cell migration.

## Discussion

Here, we show for the first time that human small intestinal organoids respond to IFNγ-treatment by undergoing transcriptional reprogramming that can subsequently promote CXCL11-driven T cell migration. IFNγ-treated small intestinal organoids undergo transcriptional changes leading to a pro-inflammatory expression profile. Treated organoids upregulated the expression of genes encoding HLAs, *CCL22* and *CCL25* (**Figure 1c-d**) important for T cell homing in the skin^37,38^ and intestine^39^ respectively. Our data evaluating *CXCL9*, *CXCL10*, and *CXCL11* expression by human intestinal organoids corroborate findings recently obtained using an induced pluripotent stem cell model^40^. We additionally provide a characterization of the functionality of the organoid system to directly modulate T cell behavior which has not been previously investigated. In addition, the use of adult stem cell-derived organoids enables the use of a relevant human model system that does not rely on reprogramming and differentiation which itself could clearly impact cellular responses. The impact of IEC-derived CXCL9, CXCL10, and CXCL11 on T cell migration was explored, demonstrating a specific role for CXCL11 driving T cell migration towards IFNγ-treated intestinal epithelium.

To evaluate how transcriptional changes functionally impact T cell behavior, we analyzed T cell migration and T cell activation in presence of IFNγ pre-treated organoids. We observed no changes in the levels of activation or proliferation of T cells compared to untreated controls. Since in each individual experiment, T cells were isolated from different blood donors, T cells were polyclonally activated using CD3/CD28 in the presence of organoids. This provides a robust method of antigen-independent stimulation. We observed that CD4^+^ and CD8^+^ T cells, regardless of their activation status, migrate towards IFNγ-treated organoids. This might be an interesting finding regarding intestinal inflammatory disease pathogenesis. In IBD and coeliac disease, T cells localize to the intestinal epithelium^18,41^. Additionally, it has been reported that T cells localize in the crypt surroundings in coeliac disease^18^ and after HSCT, relevant for GVHD pathogenesis^12,13^.

Using an antibody-blocking strategy, we observed that CXCL11, but not CXCL9 and CXCL10, drives the migration of T cells towards intestinal organoids. This finding highlights the possibility to differentially target each chemokine to modulate T cell trafficking even before allogeneic T cell reactions might emerge. In this context, CXCL10 has been reported as a biomarker of GVHD, and its expression was observed in the skin, another target organ in the disease^42^, and CXCL9-CXCL10/CXCR3 was confirmed to be critical for T cell recruitment in the lung and liver^43^. A possible explanation for the specific role of CXCL11 could be attributed to the higher affinity of CXCL11 for CXCR3 compared to CXCL9 and CXCL10^44^. Additionally, each CXCR3 ligand induces distinct conformational changes in the receptor, leading to divergent downstream effects^45,46^, thus possibly explaining our findings.

GVHD has a complex etiopathogenesis, and the role of IFNγ in its onset depends on the phase of development of the disease and the specific target tissue. In murine GVHD models, CXCL9, CXCL10 and CXCL11 expression has been found to be different among target tissues at different timepoints^47^. CXCL11 expression in the gut was high at week one after transplantation, decreased but was maintained significantly higher than controls at week two, and increased again at week three. Our ex-*vivo* data further supports a link between IFNγ-mediated epithelia damage and chemokine release that can subsequently result in further lymphocyte recruitment potentially amplifying the initial damage response.

Expression of CXCL11-related chemokines are associated with a variety of intestinal immune pathologies. For example, perfusate levels of CXCL9^48^ and serum levels of CXCL10^49^ and CXCL11^49,50^ have been reported to be higher in patients with IBD compared to healthy controls^50^. Thus far, only CXCL10 blockade has been tested in clinical trials in IBD patients, only proving to have limited effects with no remission^51,52^. Regarding coeliac disease, it has been showed that CXCL10 serum levels are increased in untreated patients, while CXCL11 was detected to be higher in duodenal biopsies at the mRNA level^53^. Currently CXCL11 has not been selected as a target for clinical intervention, but our data suggests that CXCL11 blockade may be a potential novel target for clinical intervention for preventing T cell trafficking under specific conditions.

Preventing T cell trafficking to the gut is beneficial for GVHD prophylaxis, and it is already being investigated using vedolizumab, a monoclonal antibody targeting integrin α4β7 and therefore inhibiting its interaction with mucosal addressin cell adhesion molecule 1 (MADCAM1) expressed by intestinal endothelial cells^54^. For a possible therapeutic intervention targeting CXCL11, it is important to consider that its expression might be limited to specific time frames during intestinal inflammation and disrupting the CXCL11/CXCR3 axis may only be necessary for a relatively brief period in such a therapeutic approach. In this study, we show a rapid expression of CXCL11 by organoids treated exposed to IFNγ. This suggests that a candidate moment to prevent T cell migration by targeting CXCL11 would be in the initial phases of inflammation. Indeed, organoids generated from IBD patients do not express higher CXCL11 than controls, suggesting that the upregulation of this chemokine is reversible outside of the inflammatory microenvironment even from patients exposed to long-term immune-mediated intestinal injury^55^. An important consideration is that T regulatory (T_reg_) cells counteract pro-inflammatory stimuli and can also traffic through chemokine gradients sensed by CXCR3^56^. Preventing T_reg_ cells from trafficking to the gut would eliminate a crucial mechanism for inducing tolerance, leading to increased pathogenic immune responses^57^. Additionally, the preventing T cell trafficking might impact immune surveillance in target tissues. This should also be examined in details as it has been done for vedoluzimab^58^ to evaluate risks and benefits for each patient group, Lastly, an aspect to consider in allo-HSCT patients is the Graft-versus-Tumor (GVT) effect, as many of those patients receive the transplantation to treat hematological malignancies and ideal treatment should not weaken this effect. However, it has been demonstrated that both IFNγR- and CXCR3-lacking donor cells still provide a strong GVT effect, while reducing GVHD and T cell trafficking especially in the gastrointestinal tract^56^. Collectively, those studies pose the basis of specific CXCR3/chemokine axis modulation at specific time points following transplantation to reduce GVHD while maintaining the beneficial GVT effect.

In conclusion, we have shown for the first time that human IECs exposed to IFNγ can directly modulate T cell behavior in the absence of ither cellular mediators and through the establishment of a chemoattractant gradient driven by CXCL11. We propose a potential role for CXCL11 in intestinal pathologies driven by T cell responses by recruiting CXCR3^+^ T cells to the inflamed intestine. Overall, this study supports the development of novel therapeutic strategies aimed at modulating CXCL11 activity and thereby impacting tissue-dependent IFNγ responses.

## Acknowledgments

We would like to thank all members of the Lindemans and Coffer groups for helpful discussions. Sabine Middendorp for generation of organoid lines. We thank the Hubrecht Institute FACS facility, the Olink facility at the UMC Utrecht, Noortje van den Dungen of the Epigenetics facility of the UMC Utrecht, USEQ, and Richard Volckmann from R2-support.

## Author Contributions

AC wrote the manuscript. AC and SAJ designed, performed, and analyzed experiments, and FP performed experiments. MM assisted with RNA-sequencing analysis. EM assisted with experimental design. PJC and CAL supervised the research, aided with experimental design, and wrote the manuscript. All authors contributed to the manuscript.

## Suppl. Figure legends

**Suppl. Figure 1.**
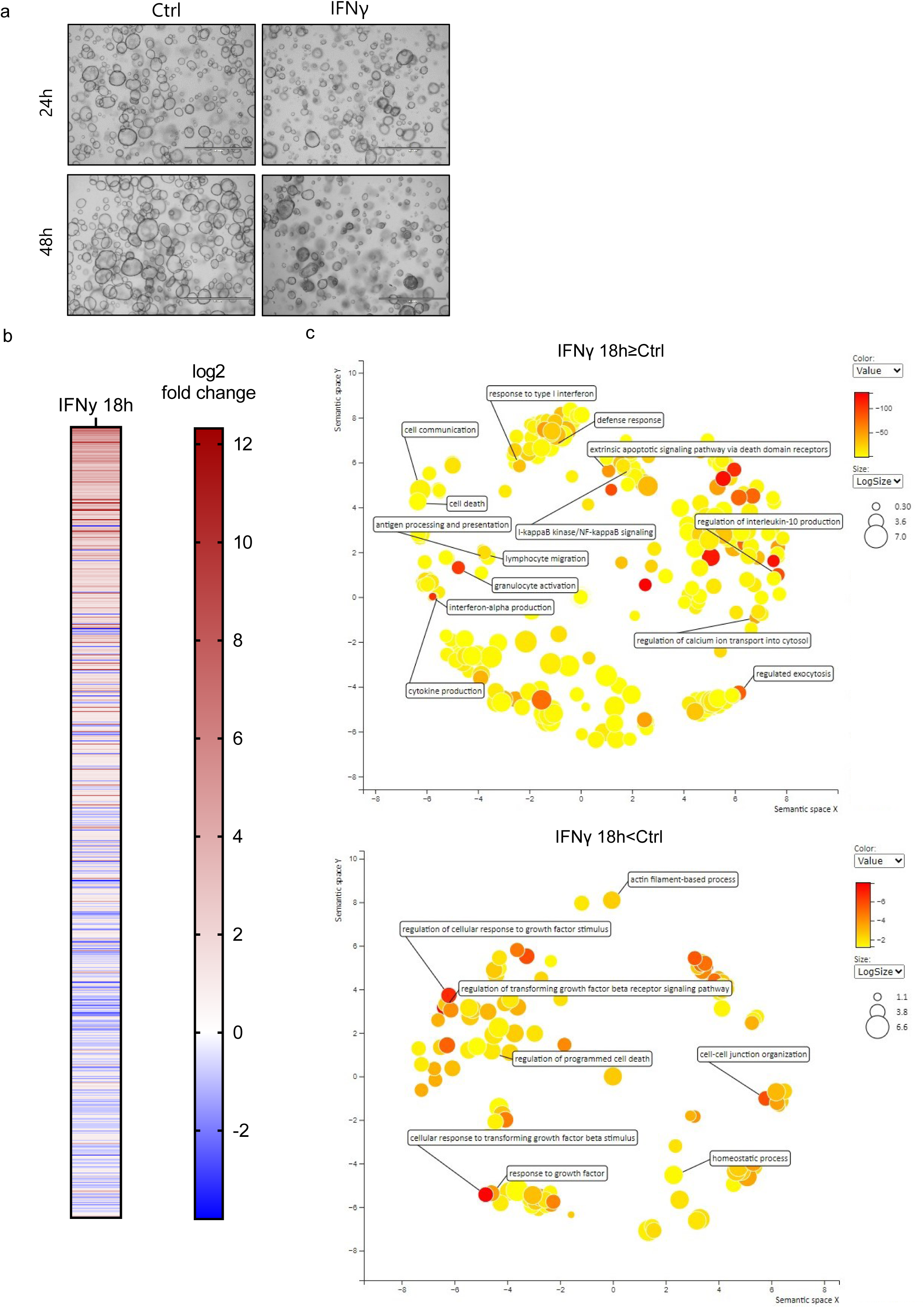
**(a)** Representative EVOS images of organoids treated for 24h and 48h with IFNγ (10 ng/ml), scale bar = 1000 μm. **(b)** Heatmaps of all significant DEGs after treatment with IFNγ for 18h compared to untreated controls (DESeq2, padj<0.05), ordered per log2FC (higher to lower). **(c)** REVIGO analysis of GO gene sets up- or downregulated compared to untreated control organoids obtained from R2 analysis (padj<0.05).

**Suppl. Figure 2.**
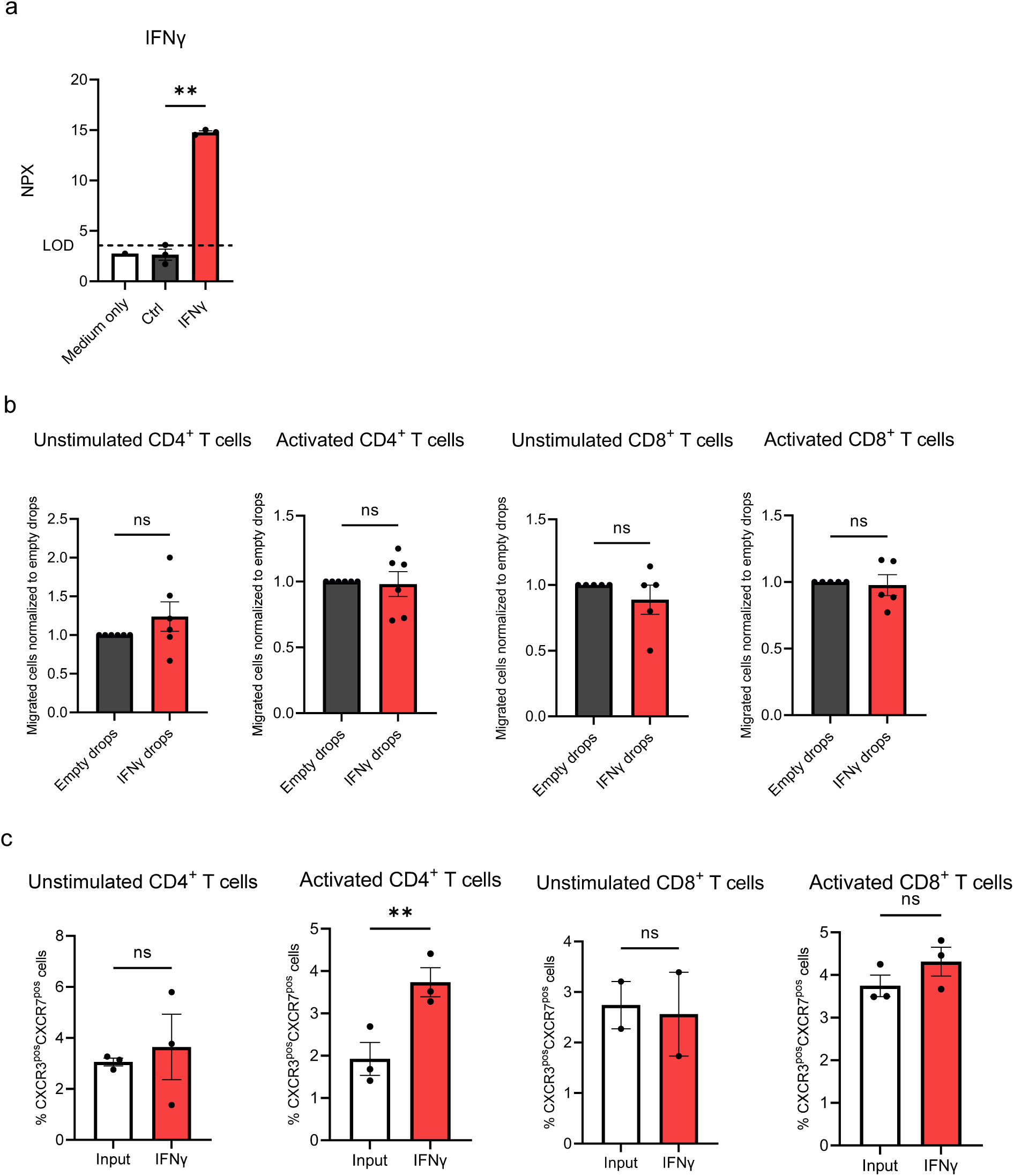
**(a)** IFNγ levels detected by Olink proteomics. Each data point indicates the levels of the detected protein in the conditioned media from each organoid condition (log2 scale), mean with SEM, ANOVA. **(b)** Normalized number of unstimulated and pre-activated CD4^+^ and CD8^+^ T cells that have migrated overnight from a 3 μm-pore sized insert (upper compartment) to the lower compartment containing empty Matrigel drops (without organoids) that were treated with IFNγ (10 ng/ml) for 48h, and then refreshed for 24h prior to the beginning of the assay, counted by FACS. n≥5 T cell donors with 1 organoid line, each data point indicates a T cell donor, mean with SEM, Student *t* test. **(c)** (Membrane) CXCR3 and CXCR7 of unstimulated and pre-activated CD4^+^ and CD8^+^ T cells that have migrated overnight from a 3 μm-pore sized insert (upper compartment) to the lower compartment containing organoids that were treated with IFNγ (10 ng/ml) for 48h, and then refreshed for 24h prior to the beginning of the assay. % of CXCR3 and CXCR7 double positive cells are compared to the percentage in the input material, which consists of T cells cultured for the whole duration of the migration assay in a well containing empty Matrigel drops (no organoids). N≥2 T cell donors with 1 organoid line, each data point indicates a T cell donor, mean with SEM, Student *t* test.

